# A high-resolution pipeline for 16S-sequencing identifies bacterial strains in human microbiome

**DOI:** 10.1101/565572

**Authors:** Igor Segota, Tao Long

## Abstract

We developed a <u>Hi</u>gh-resolution <u>M</u>icrobial <u>A</u>nalysis <u>P</u>ipeline (HiMAP) for 16S amplicon sequencing data analysis, aiming at bacterial species or strain-level identification from human microbiome to enable experimental validation for causal effects of the associated bacterial strains on health and diseases. HiMAP achieved higher accuracy in identifying species in human microbiome mock community than other pipelines. HiMAP identified majority of the species, with strain-level resolution wherever possible, as detected by whole genome shotgun sequencing using MetaPhlAn2 and reported comparable relative abundances. HiMAP is an open-source R package available at https://github.com/taolonglab/himap.

---

Recent studies have demonstrated the importance of human gut microbiome in contribution to many diseases, as well as anti-tumor effects in cancer immunotherapies. Despite their progress, a major open question encompassing many clinical studies, is the identity of causal gut bacterial species or strains. The most prevalent method for microbiome investigation is 16S ribosomal RNA (rRNA) amplicon sequencing, in which 16S rRNA gene hypervariable regions are PCR amplified and sequenced. Common analysis pipelines^1,2^ coarse-grain sequencing reads into 97% similarity clusters called OTUs (Operational Taxonomic Units) and use a reference database to identify bacteria typically at family or genus levels. Even though this approach has demonstrated success in exploratory studies^3,4^, it presents a barrier for clinical studies where the goal is to identify specific species or strains associated with the phenotype of interest, such as immunotherapy responsiveness^5^.

The majority of clinically relevant strains from human gut have being sequenced. However, this information is presently underutilized in reference 16S databases. Major 16S databases are RDP DB^6^, Greengenes^7^, and SILVA^8^. Each of these databases contain millions of sequences and have taxonomy computationally assigned, but only a few percent of these sequences represent named strain isolates. Greengenes and SILVA impede reproducibility due to unavailability of the 16S sequence sets used for constructing phylogenetic tree for taxonomy assignment. Only SILVA is updated every year and, out of the three databases, contains the most unique species and strains (**Supplementary Figure 1 and Note 1**). Many identical full-length 16S sequences are assigned different names in different databases (**Supplementary Figure 2**), suggesting high uncertainty of computational assignments^9^.

Recently, independent research groups have developed various denoising algorithms to infer exact sequences by accounting for sequencing errors, as an alternative to the 97% OTU approach^10–12^. A major issue is that denoised sequences are still matched against a small subset of aforementioned databases (e.g. Greengenes clustered at 97% identity), potentially obfuscating the biggest advantage of exact sequence inference. At present, more than 100,000 bacterial and archaeal strains have sequenced genomes, with the number doubling every 1.3 years (**Supplementary Figure 3**). Many of these strains come from the human gut microbiome and exact matches to these sequenced strains can be far more useful for functional investigation than a coarse and uncertain taxonomic assignment. Furthermore, present pipelines underutilize or ignore the information of copy numbers and sequence variations of 16S rRNA genes in fully sequenced genomes, further contributing to the inaccuracies of species identification and abundance estimate (**Supplementary Figure 4**).

To address these issues, we developed the R package HiMAP as a bioinformatics pipeline for processing 16S sequencing data of human microbiome, from raw FASTQ reads to the final table with estimated abundances of identified strain groups. HiMAP implements a *de novo* 16S database, obtained by automatically extracting and curating bacterial and archaeal 16S sequences from NCBI Genome and RefSeq databases (**Figure 1**). HiMAP database contains more unique species and strains than any major database (**Supplementary Figures 1 and 5**), is fully documented, open source (https://github.com/taolonglab/himapdb), and can be updated at any time. HiMAP database consists of a standardized FASTA and a tab-delimited text file, thus can be easily further curated when needed.

**Figure 1.**
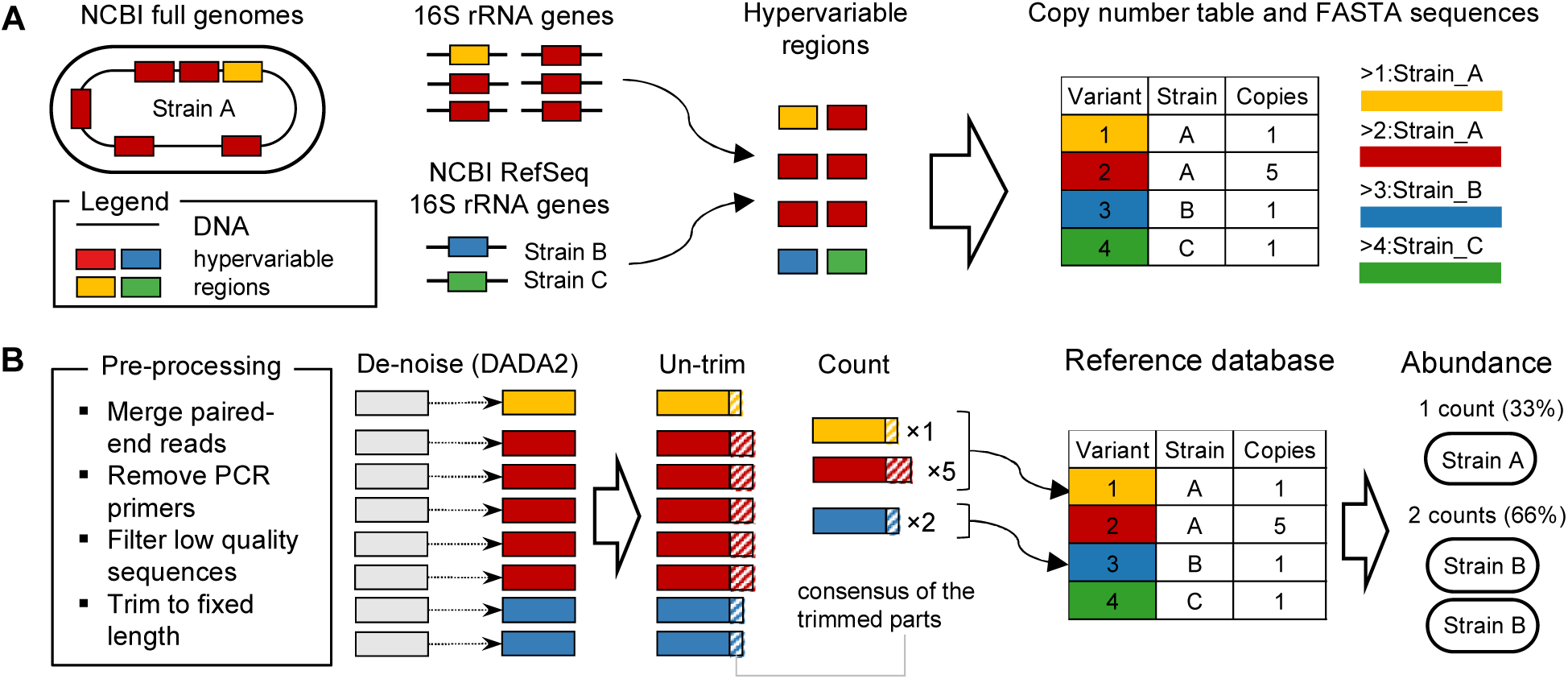
HiMAP pipeline description. **A.** HiMAP reference database is constructed using (i) 16S rRNA gene sequences from NCBI Genome database (Bacteria and Archaea) and (ii) NCBI RefSeq 16S sequences. Using standard PCR primer sequences, we extract specific hypervariable regions (e.g. V3-V4) from all full or partial 16S sequences. For each strain, we then obtained the number of unique hypervariable region variants. **B.** Pipeline begins by pre-processing demultiplexed paired-end reads in FASTQ files: read merging, PCR primer removal, quality filtering and fixed-length trimming. We use DADA2 to denoise sequences and then concatenate the consensus sequence of the trimmed region back, to improve database matching. These sequences are counted for each sample and aligned against HiMAP reference database. Finally, we assign a count for each set of best-match strains (an Operational Strain Unit; OSU) using a linear model using all detected variants of each strain.

HiMAP workflow (**Figure 1** and Online Methods) starts by pre-processing: paired-end reads are merged, PCR primers are removed and sequences are quality filtered and trimmed to fixed-length. Trimming length is automatically determined, based on the 99% quantile of sequence lengths (trimming length is a parameter that can be changed). Denoising is then performed using DADA2^11^, but with sample-dependent and Bonferroni-adjusted p-value threshold (OMEGA_A) of 10^−4^. After denoising, the consensus sequences of trimmed regions are concatenated back (un-trim), sequences are counted, and chimeric reads are removed. The resulting sequences that differ only in shift or length are pooled using an extremely fast HiMAP *collapse* function.

Next, collapsed sequences are aligned to the HiMAP database and only alignments with the highest score are kept. For data from human gut microbiome studies, multiple 100% database matches are common, so we introduce the concept of an Operational Strain Unit (OSU). An OSU is a set of strains impossible to distinguish by the sequenced hypervariable region, and consist of either a single shared sequence or multiple intra-strain sequence variants. HiMAP uses all the information of copy numbers and sequence variations of the hypervariable region of 16S rRNA genes in the genomes to pin down the strains in these OSUs, by solving a linear model that best explains the observed sequence counts with estimated OSU counts. For a sequence with less than 100% database match, an OSU is a set of strains with the most similar sequences. The final output consists of a table with counts of each OSU for each sample, with NCBI assigned taxonomy.

We tested HiMAP on three data sets: (i) Human Microbiome Project (HMP) 20-species mock community with staggered 16S gene concentrations (V3-V4 region, 1 sample)^13^, (ii) a healthy human sample with matching whole genome shotgun (WGS) sequencing data (V3-V4 region, 1 sample)^14^ and (iii) DIABIMMUNE longitudinal study of infants with matching WGS data (V4 region, 780 samples)^15^. We analyzed WGS data using a marker-gene based pipeline MetaPhlAn2^16^ and used the results as the reference for comparing various16S pipelines including HiMAP.

In the HMP mock community, HiMAP identified all 20 species at 100% identity. The OSU abundances of the 20 species follow the cell abundances well (**Figure 2A, Supplementary Table 1**), with part of the uncertainty caused by the difficulty to infer the exact 16S gene copy number when there are multiple matching strains (**Supplementary Figure 6**). 13 species have been uniquely identified, while for 7 an output is typically a short list (2-10) of exactly matched species. Strain-level resolution (fewer than 10 strains) is challenging, because many strains share an identical hypervariable region, but sometimes is possible (**Figure 2B**). For *A. odontolyticus* and *D. radiodurans* this is likely due to having only 6 and 3 (respectively) strains per species in the HiMAP database. For *S. epidermidis, C. beijerinckii* and *B. vulgatus* (456, 26 and 22 database strains), strain-level resolution was achieved by identifying multiple hypervariable region variants, present at the predicted ratios (**Supplementary Figure 7**).

**Figure 2.**
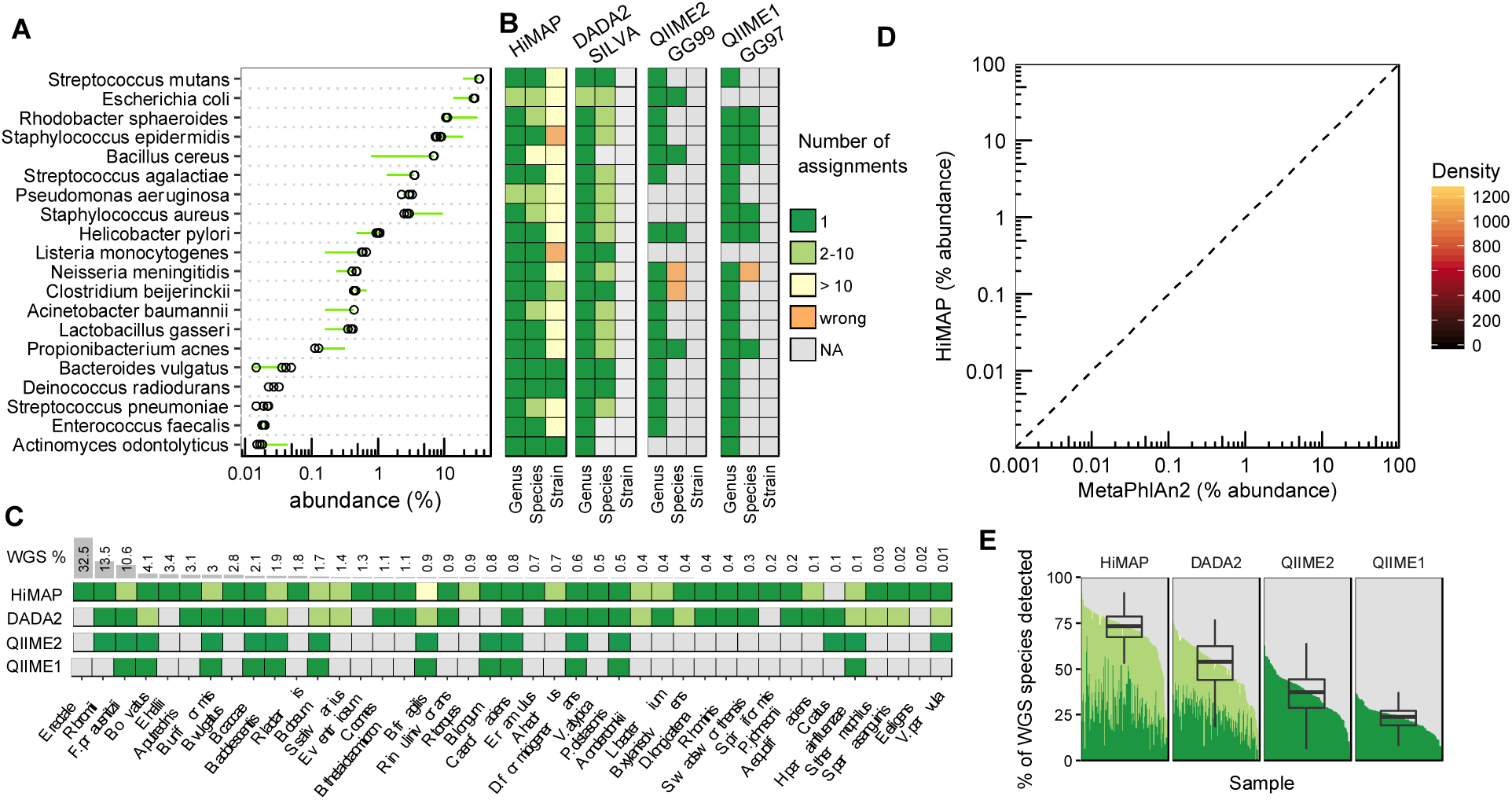
Identification and abundances of strains of the HMP mock community, V3-V4 region (A-B), healthy human gut microbiome, V3-V4 region (C) and DIABIMMUNE human gut microbiome samples, V4 region (D-E), where both human gut microbiome samples have matching WGS data. **A.** Abundances of each mock community strains as estimated by HiMAP (open circles) and as calculated from listed reference 16S rRNA gene abundances (green circles). Green line segments show the (log) difference between estimate and the reference abundance. **B.** Identification of genus, species and strain ranks by HiMAP, DADA2, QIIME2 and QIIME1 pipelines. **C**. Identification of 41 bacterial species obtained from MetaPhlAn2. **D.** Comparison of abundances estimated by HiMAP (y axis) and MetaPhlAn2 (x axis) for each matching species or strain. **E.** Percentage of WGS species identified by each pipeline. On average (median), HiMAP identified 73% of WGS species, DADA2 SILVA 54% and QIIME2 GG99 37%. Only 16S samples with more than 100,000 reads are shown.

For comparison, we also tested three other pipelines: DADA2 (SILVA NR v132 database), QIIME2 and QIIME1 (Greengenes v13.8 clustered at 99% and 97% identities, abbreviated GG99 and GG97), following recommended workflows (**Supplementary Note 3**). DADA2 identified 17 species, QIIME2 identified 6 (2 incorrectly assigned: *N. meningitidis* as *N. cinerea* and *C. berijerinckii* as *C. butyricum*; also in ^12^), while QIIME1 identified 7, including *S. aureus* that QIIME2 missed (**Figure 2B)**. HiMAP output on average 8 additional OSUs at low abundances which include false positives, contaminations and denoising artifacts, comparable to DADA2 and QIIME2, while QIIME1 output 160 additional OTUs, occurring across replicates and spanning the entire abundance range (**Supplementary Figure 8**).

Next, we analyzed a healthy human gut microbiome sample with matching 16S and WGS data (1.2M reads each)^14^. MetaPhlAn2 reports *Eubacterium rectale* as the most abundant (32.5%) species (**Figure 2C**), consistent with results reported by HiMAP, with comparable abundance (19%) and 100% identity. However, none of the other pipelines output even the *Eubacterium* genus. We confirmed this species’ presence in the 16S data by BLAST-ing the most abundant sequence (after read merging and PCR primer removal) against the NCBI 16S rRNA database. The single best scoring alignment is an exact full-length match to *E. rectale* (**Supplementary Figure 9**).

HiMAP identified 40 out of 41 MetaPhlAn2 species, belonging to OSUs with ≥ 97% identity to HiMAP database (MetaPhlAn2 assignments are not based on 100% identity to marker-genes either). HiMAP and MetaPhlAn2 both identified *Lachnospiraceae bacterium* 5.1.63 FAA on a strain level, while a single missing species, *Coprococcus catus*, is not in the HiMAP database. Its sequence is assigned *Lachnoclostridium* spp. (96% identity), but manual BLAST search against the NCBI Nucleotide database, reveals an exact match to the *Coprococcus catus* GD/7 draft genome. Overall, HiMAP shows excellent agreement with MetaPhlAn2 analysis, identifying essentially all of its species while capturing a much higher bacterial diversity: 440 OSUs vs 47 MetaPhlAn2 assignments (6 had “unclassified” species).

Finally, on a DIABIMMUNE dataset, WGS was done at ∽500 fold higher sequencing depth than 16S (**Supplementary Figure 10**). We compared abundances for all matching species or strains, by re-normalizing them to 100% in each sample. The correlation in abundances between HiMAP 16S and MetaPhlan2 WGS analysis is very good (Pearson R = 0.72, **Figure 2D**), with ± 3 fold deviations, comparable to the mock community abundance uncertainty. On 16S samples with ≥ 10^5^ reads, HiMAP recovered on average (median) 73% of strains or species from MetaPhlAn2, while DADA2, QIIME2 and QIIME1 pipelines recovered on average 54%, 37% and 24% (**Figure 2E**) and missed many low abundance species (**Supplementary Figure 11**).

Half of all HiMAP OSUs have a 100% match to one or more species in the database (75% OSUs have ≥ 99% match; **Supplementary Figure 12**), demonstrating the advantage of cross-referencing exact sequences with the HiMAP database when the database coverage is good, as is the case for human microbiome studies. As more strains are being sequenced, HiMAP database can be expanded to also improve other uses cases, such as the mouse microbiome studies. Furthermore, the full genome information can be better utilized for functional predictions^17^.

## Online Methods

### Database construction

HiMAP reference database is generated by a series of Python scripts, with the detailed instructions and source code available at https://github.com/taolonglab/himapdb. First, we use *dl*_*genome*_*meta.py* to download bacterial and archaeal genome assembly summary tables from the NCBI Genomes database FTP server. The same script generates cleaned-up strain names used throughout the pipeline, and for each unique strain name selects only one genome assembly. This assembly is selected by prioritizing first (i) an assembly level (Complete Genome > Chromosome > Scaffold > Contig), then (ii) RefSeq category (reference genome > representative genome > no annotation) and then (iii) release date, by taking a newer assembly in case of ties in (i) and (ii). In case there are still multiple assemblies tied, we select the first one from the list. The output from this script are two files for each kingdom (Bacteria and Archaea) with direct links to FASTA sequences and GFF annotations for each genome assembly. Then, *download.py* downloads these sequences and annotations. This is the most time-consuming step as the download consists of about 50 GB of compressed sequence data. Next, *extract.py* extracts annotated 16S sequences from the FASTA files, based on the annotation in the corresponding GFF files, and stores all sequences in a single FASTA file. We then generate a reduced sequences that correspond to the hypervariable region amplified by a specific set of forward and reverse PCR primers. The script *count.py* takes as input PCR primers and the FASTA file with 16S sequences. For each strain, it extracts and counts the number of unique sequences for a specific hypervariable region. This is done using the command line version of nucleotide BLAST (blastn) to find the regions between forward and reverse primers. It outputs a FASTA file with sequences and a table with assembly identifiers, strain names and counts.

Next, the script *dl*_*refseq.py* uses NCBI web eUtils to download search results from the RefSeq database which match the search string “16s ribosomal RNA[Title] NOT uncultured[Title] AND (bacteria[Filter] OR archaea[Filter]) AND (1000[SLEN] : 2000[SLEN]) AND refseq[filter]” and saves the sequences to a FASTA file. Finally *add*_*refseq.py* adds the information from this FASTA file to the FASTA file and tables from full genome assembly, for a specific primer set and for each new strain.

Final filtering step is performed by an R script *remove*_*taxonomic*_*outliers. R*, by excluding “taxonomic outliers” or “duplicate species”. If, within a set of strains sharing the exact 16S hypervariable region, 99% or more strains share a single taxonomic rank (Family, Order, Class or Phylum), the remaining ≤ 1% strains are removed. In addition, strains with double species annotations, such as “Psychrobacter immobilis Neisseria meningitidis” are removed as well. These excluded are saved in a separate table for each hypervariable region, e.g. *excluded*_*strains*_*from*_*himap*_*database*_*v4.txt*. and consist of 87 and 180 strains for V3-V4 and V4 hypervariable region, respectively.

### FASTQ preprocessing

The raw (demultiplexed) FASTQ data is pre-processed in three steps: (i) paired-end read merging, (ii) PCR primer removal and (iii) quality filtering and fixed-length trimming. Read merging is performed by a HiMAP function *merge*_*pairs* which merges reads using an overlap version of the Needleman-Wunsch alignment algorithm and calculates the correct posterior error probabilities and corresponding phred scores of the overlapping sequence region^18^. Since matching base-calls in the overlap region can dramatically increase phred scores, this allows more accurate identification of correct sequences later in the pipeline. PCR primers are removed with a HiMAP function *remove*_*pcr*_*primers*, which uses fitting alignment version of the Needleman-Wunsch alignment. Quality filtering is performed with a HiMAP function *filter*_*and*_*trim*, a wrapper for *filerAndTrim* function from the DADA2 package. By default, reads are discarded if their sequence contains any Ns, they have more than 2 expected errors, are shorter than 20 nts, if they match phiX genome or are shorter than a fixed-length threshold. This (user-settable) threshold is determined by calculating a 1% quantile length of a sample of sequences after PCR primer removal, such that 99% of the sequences are above this threshold.

### Denoising

Denoising is performed using a function *dada*_*denoise*. Internally, this function runs in three steps. First, error rates are learned using *dada2::learnErrors* function, based on the pooled set of a number of samples that have at least 1 million reads combined. Second, these error rates are used to run DADA2 algorithm on each sample separately to obtain a list of denoised sequences. Briefly, the DADA2 algorithm partitions similar sequences around a central sequence, estimated to be the true sequence of each partition, based on their abundance and quality scores. The key step of the partitioning process is assignment of a sequence to either a new or a pre-existing partition. This is done by calculating a probability (based on the Poisson model) of observing at least an observed number of identical reads of that sequence with the average quality score profile. If this probability is lower than a specified threshold OMEGA_A, then the sequence is assigned to a new partition. The DADA2 default (sample independent) value for OMEGA_A = 10^−40^, which we changed to sample-specific value of 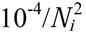 where *N*_*i*_ is the number of unique reads in the sample *i*. This is a Bonferroni-adjusted p-value of 10^−4^, where we use 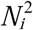 as the estimate for the number of pairwise comparisons performed by the DADA2 partitioning algorithm. Finally, in the last step full length sequences (before fixed-length trimming) are retrieved by backtracking to the pre-trimmed FASTQ file, for the center sequence of each partition. The consensus sequence of the trimmed part is then concatenated to the trimmed sequence. For example, for V3-V4 region the concatenated part is around 20 nt long.

### Sequence abundances

After denoising, abundance of each sequence in each sample is calculated using a function *sequence*_*abundance*. First, central sequences from each DADA2 partition are assigned read counts by adding up the counts of each unique sequence in that partition. Second, chimeric reads are removed using a DADA2 function *removeBimeraDenovo* and sequences from all samples are checked for sequences that differ only in shift or length. This part is done using an optimized BLAST alignment, through a HiMAP *collapse* function. This alignment uses very large word size (200-400 nt, based on the sequence lengths) and runs extremely fast even with large number of samples and sequences.

### Matching sequences against a reference database

The final list of sequences is matched against a HiMAP reference database for a specific hypervariable region, using a MegaBLAST from the *blastn* command line tool (bundled with the HiMAP package). User can easily provide their own or a third-party reference database at this step. For each sequence, the best BLAST hits are saved, based on the alignment score (using parameters: match = 5, mismatch = −4, gap open = −8, gap extend = −6), with a minimum percentage identity of 75% and word size of 50 nt. HiMAP provides pre-generated databases for V4 and V3V4 hyper-variable regions. For exact (100%) matches to the database, copy number information of each hyper-variable region is retrieved from a separate table.

### Abundance estimation of Operational Strain Units (OSUs)

The abundance estimation of OSUs is split into two separate parts. Sequences with < 100% matches to the database are each assigned a single OSU with raw count equal to the sequence raw count. Exact 100% matches to database strains are processed in several steps. Each OSU is a bin of strains sharing exact same sequence variant spectrum, with potentially different copy numbers. One OSU bin could contain a single sequence variant *v*, with 3 different strains having copy numbers 1, 3 and 5. In that case the copy number of that OSU bin is assigned an integer value of an average copy number over all strains (3), so the OSU spectrum is {*v*:3} Another OSU bin may contain multiple sequence variants, e.g. *v*_*1*_ and *v*_*2*_, with copy numbers 5 and 1: {*v*_1_:5, *v*_2_:1}.

Since multiple strains (and OSUs) can share the exact same sequence variants, the abundances of each OSU are estimated using a linear model, as follows. The abundance *s*_*i*_ of each sequence variant *i* is obtained by summing up contributions *a*_*i*j_ from each OSU, *j:* 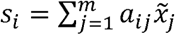, where *a_ij_* is the copy number of sequence variant *i* in the OSU *j, m* is the total number of OSUs in the database and 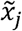 is the real (unknown) count of that OSU and *i* ∈ {1, …, x*n*} with *n* being the total number of denoised sequences in a specific sample. This system of equations can be represented in matrix form as 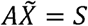, e.g. for n = 3 and m = 2:

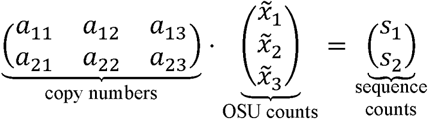

where each column of the A matrix contains the copy number of each OSU for each detected sequence (its spectrum). Due to the uncertainty in the sequence counts *s*_*i*_, in all practical cases this system is overdetermined and does not have an exact solution. However, an estimate for the exact solution can be obtained by finding OSU counts *X* that minimize the square of the residuals: ‖*Ax - B*‖^2^.

The issue with this solution is that it is very prone to overfitting. For example, assume a sample contains the strain with two sequence variants with copy numbers 3 and 1, i.e. a spectrum {*v*_1_:3, *v*_2_:1} and that experimentally we obtain sequence counts of 279 and 102 for these two variants, respectively. It is often the case that a database contains strains that have only variant *v*_*1*_ or *v*_*2*_. In this case the optimal solution would have count 279/3 = 93 for the strain with {*v*_1_:3, *v*_2_:1}spectrum and 102-93 = 9 for the strain with {*v*_2_:1} spectrum. This is an example of overfitting due to the uncertainty in counts. To address this, we added a regularization term to the minimization function:

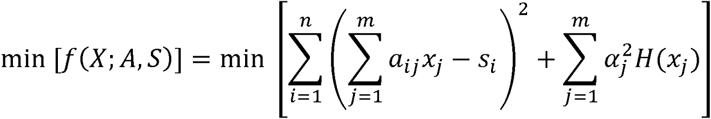

The first term in the brackets is the sum of all the squared residuals ‖*Ax - B*‖^2^, while the second regularization term adds a *a*_*j*_ cost to the introduction of each new OSU *j*. The *H*(*x*) is a Heaviside step function. We solve this minimization problem in two steps, to speed up computation. First, the solution is obtained for the non-regularized function using a function *lsei* from an R package limSolve ^19^, with constrains *x*_*j*_ ≥ 0 for all *j*. This part eliminates large number of missing OSUs, and therefore large number of columns from the A matrix. To speed up the minimization in the next step we transform the reduced A matrix into a block-matrix form and perform minimization on each independent block separately. The block-matrix transformation is done by converting the rows and columns of matrix A into an undirected graph, using an R package igraph^20^. In this graph, each row is connected to a column, if the copy number in that matrix field is greater than zero. Then, all the connected components of the graph are obtained using a function *components* and each of these components are used to extract block-matrices of A. Each block is optimized separately by setting the regularization weights *a*_*j*_ to the un-regularized solution is *x*_*j*_ Then, the regularization term has the form 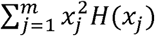. The regularized solution is obtained using *pso* function from the R package pso (Particle Swarm Optimizer)^21^ with parameters maxit = 10 and vectorize = TRUE. This function is re-initialized and run 1000 times (can be changed using option *pso*_*n*) and the solution with a set OSUs is selected that has the lowest value of function *f*.

Finally, the OSUs and counts with exact matches are combined with the inexact matches to show the final table with counts for each OSU and for each sample.

### Taxonomic assignment

HiMAP can assign taxonomy to each OSU, based on the taxonomy from each species in that OSU and its assignment on the NCBI Taxonomy database. This is done by downloading a NCBI Taxonomy database dump (ftp://ftp.ncbi.nlm.nih.gov/pub/taxonomy/), converting it to a tab-delimited table using a Python script *ncbitax2lin.py* (from https://github.com/zyxue/ncbitax2lin) and the resulting table is bundled with the HiMAP package and is added to the OSU table using the HiMAP *taxonomy()* function.

