## Supplementary Information for "A high-resolution pipeline for 16S-sequencing identifies bacterial strains in human microbiome"

Igor Segota & Tao Long
Sanford Burnham Prebys Medical Discovery Institute, 10901 North Torrey Pines Road, La Jolla, CA 92037, USA

### Supplementary Figures

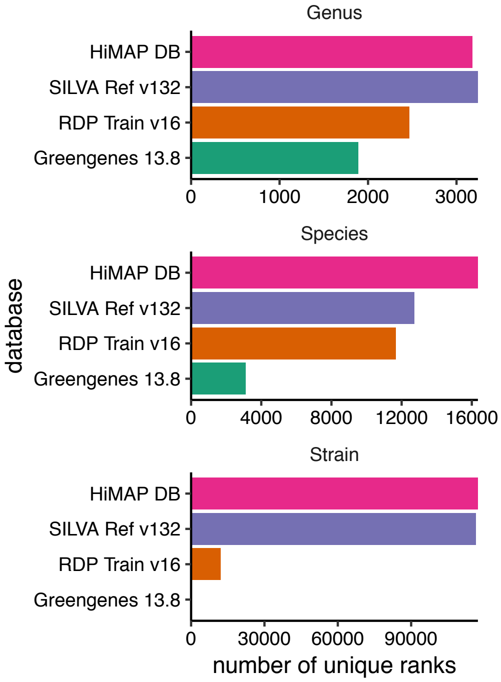

Figure 1. Number of unique genera, species and strains in the HiMAP database, Greengenes 13.8 database clustered at 99% identity, SILVA Reference v132 database and RDP DB v16 training set.

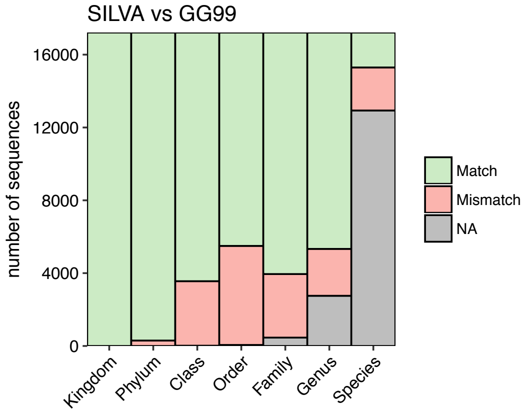

Figure 2. Comparison of taxonomic assignment for exact full length 16S sequences (≥1200 nt) in SILVA and Greengenes databases. Green indicates the number of sequences having the same taxonomic rank, red indicates different rank and gray indicates that either SILVA or (more commonly) Greengenes is missing a rank assignment.

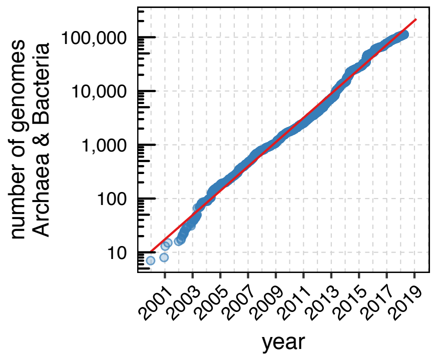

Figure 3. Number of sequenced genomes for archaea and bacteria in NCBI Genome database as a function of time. The number follows an exponential law and doubles every 1.3 years. We obtained the number of genomes for archaea and bacteria by calculating a cumulative sum of genomes present in assembly_summary.txt files at ftp://ftp.ncbi.nlm.nih.gov/genomes/refseq/bacteria/ and ftp://ftp.ncbi.nlm.nih.gov/genomes/refseq/archaea/.

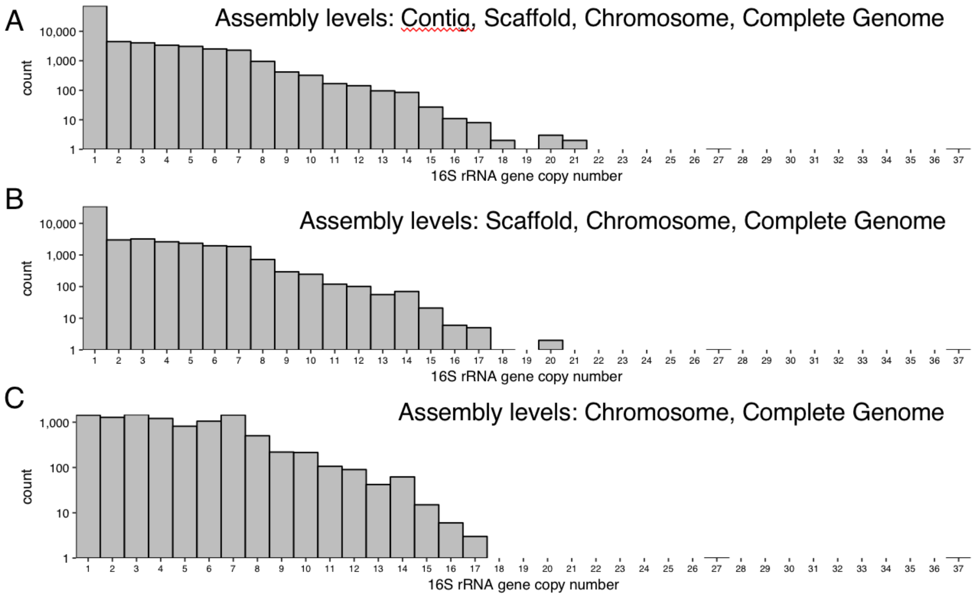

Figure 4. 16S rRNA gene copy number for strains with sequenced genomes from NCBI Genome database. 16S copy number information was obtained from the NCBI Prokaryotic annotation pipeline for each full genome assembly, while filtering out sequences shorter than 1000 and longer than 2000 nucleotides in length, then counting the number of annotations of “16S ribosomal RNA” for each unique strain. While most assemblies have one copy (A), copy number can be as large as 37. This is true especially for strains which have near-full assembly (assembly level “Scaffold”) or fully assembled genome, i.e. assembly levels “Chromosome” or “Complete Genome” (B,C).

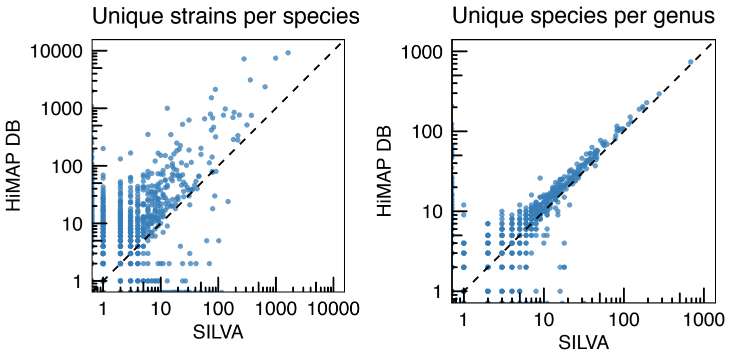

Figure 5. Comparison between HiMAP DB and SILVA, each point showing the number of unique strains per matching species (left) or the number of unique species per matching genus (right).

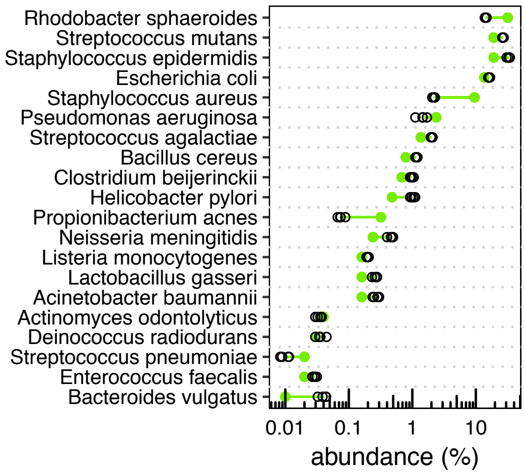

Figure 6. Species abundances as predicted by HiMAP, when a reduced V3-V4 database is used, consisting only of 20 reference strains, so each is assigned its reference 16S gene copy number. The remaining difference in abundance between the data (black open circles) and predicted values (solid green circles), is likely due to PCR amplification bias (e.g. for *P. acnes* which has 1 mismatch in the reverse 805R primer) and the uncertainty in determining precise 16S gene copy number due to effects such as growth-rate dependency of gene copy numbers due to multiple concurrent replication forks[1] or possible nonlinearities in PCR amplification[2].

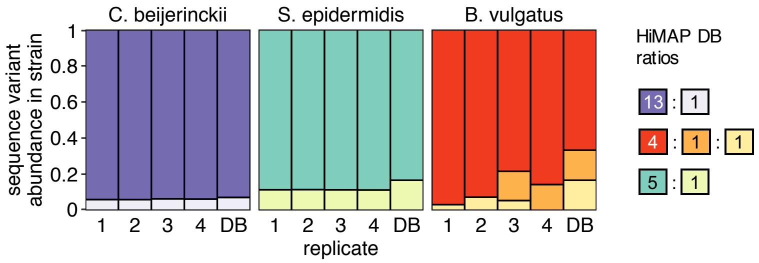

Figure 7. Strain-level resolution from the HMP mock community[3]. *Clostridium beijerinckii* NCIMB 8052 is identified as *Clostridium beijerinckii* NCIMB 8052 or *Clostridium beijerinckii* ATCC 35702 SA-1, both of which have the two exact same unique 16S rRNA gene variants, in the ratio 13:1. "DB" shows the 13:1 ratio, and the 4 other columns show this ratio in 4 technical replicates. *Staphylococcus epidermidis* FDA PCI 1200 is mis-identified as *Staphylococcus epidermidis* RP62A, *Staphylococcus epidermidis* ET-024 or *Staphylococcus epidermidis* FDAARGOS 157 because one of the sequence variants from FDA PCI 1200 is missing and these three strains do not have that variant, but have other two at a comparable ratio. *Bacteroides vulgatus* ATCC 8482 is uniquely identified in all 4 replicates, even when only 2 out 3 variants were identified.

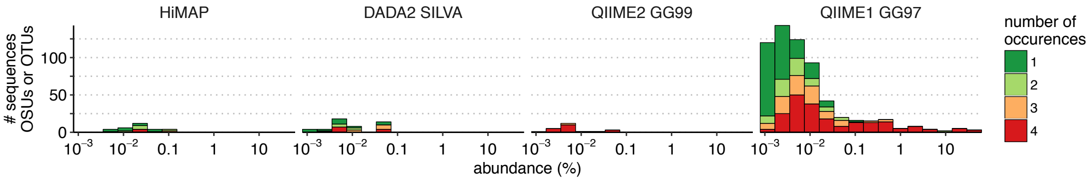

Figure 8. Distribution of false positives, contaminants, denoising and clustering artifacts in Zheng et al. 2015 HMP mock community data with 4 technical replicates. Color denotes the number of replicates an OSU, OTU or an exact sequence variant occurs in. QIIME1 produces by far the most extra OTUs, spanning an entire abundance range.

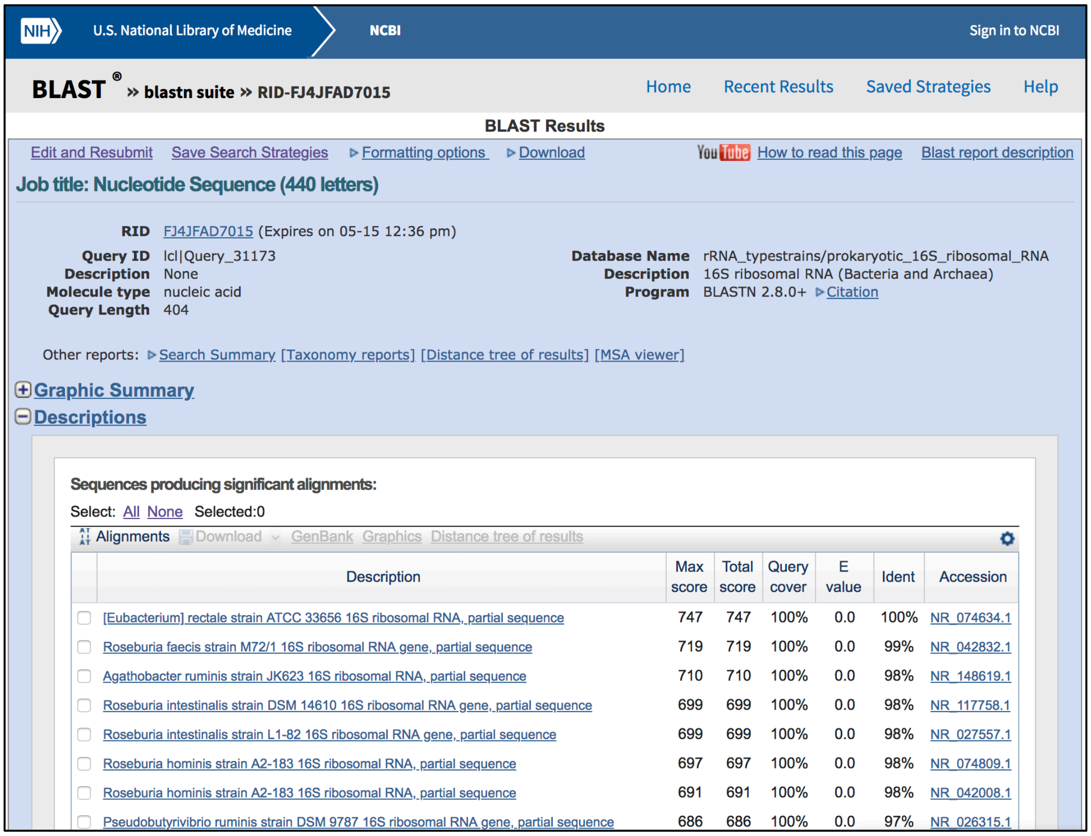

Figure 9. BLAST alignment of the most abundant sequence from Jovel et al. 2016 healthy human sample ("DON2A"), to the NCBI 16S ribosomal RNA (Bacteria and Archaea) database.

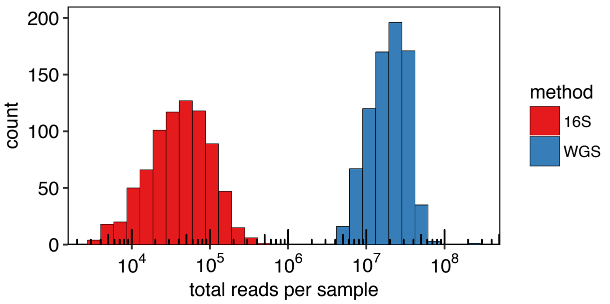

Figure 10. Sequencing depth (total number of paired-end reads in the raw FASTQ files) for 16S (red) and WGS (blue) data in the DIABIMMUNE study for 780 matching samples.

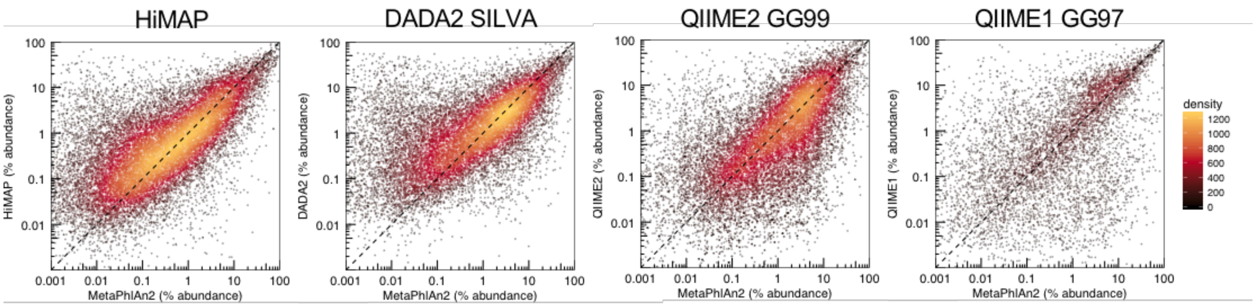

Figure 11. Comparison of abundance estimates between MetaPhlAn2 and four 16S pipelines: HiMAP, DADA2, QIIME2 and QIIME1.

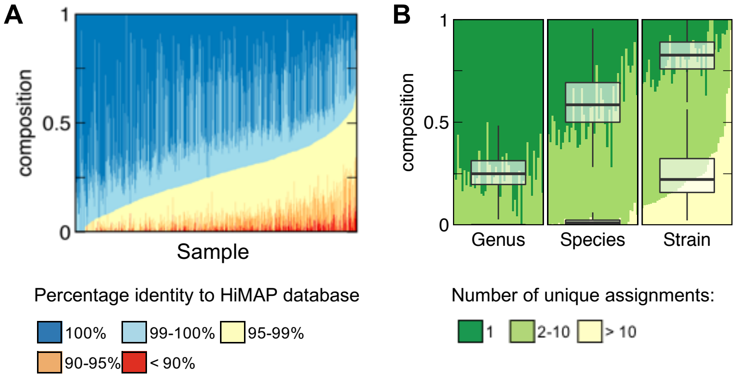

Figure 12. **A**. Composition of percentage identity of all HiMAP OSUs to the HiMAP database. About 50% OSUs have exact (100%) database matches, and about 75% have high identity (≥ 99%) matches. **B**. Composition of the number of unique assignments per taxonomic rank. About 75% of OSUs have unique genus, 45% have unique species and 18% have a unique strain assignment.

### Supplementary Tables

Table 1. Abundances of staggered HMP mock community data from Zheng et al. 2015. % cell abundance is obtained by dividing “% 16S rRNA gene abundance” by the “16S copy number” and “% normalized cell abundance” is then this number normalized to 100%.

| Species | % 16S rRNA gene abundance | Strain name in HiMAP database | 16S copy number | % cell abundance | % normalized cell abudance |
|---|---|---|---|---|---|
| Escherichia coli | 21.91 | Escherichia coli K-12 MG1655 | 7 | 3.13 | 13.63 |
| Rhodobacter sphaeroides | 21.91 | Rhodobacter sphaeroides 2.4.1 | 3 | 7.30 | 31.80 |
| Staphylococcus epidermidis | 21.91 | Staphylococcus epidermidis ATCC 12228 | 5 | 4.38 | 19.08 |
| Streptococcus mutans | 21.91 | Streptococcus mutans UA159 | 5 | 4.38 | 19.08 |
| Bacillus cereus | 2.19 | Bacillus cereus ATCC 10987 | 12 | 0.18 | 0.79 |
| Clostridium beijerinckii | 2.19 | Clostridium beijerinckii NCIMB 8052 | 14 | 0.16 | 0.68 |
| Pseudomonas aeruginosa | 2.19 | Pseudomonas aeruginosa PAO1 | 4 | 0.55 | 2.38 |
| Staphylococcus aureus | 2.19 | Staphylococcus aureus USA300 TCH1516 | 1 | 2.19 | 9.54 |
| Streptococcus agalactiae | 2.19 | Streptococcus agalactiae 2603V/R | 7 | 0.31 | 1.36 |
| Acinetobacter baumannii | 0.219 | Acinetobacter baumannii ATCC 17978 | 6 | 0.04 | 0.16 |

| | | | | | |
|---|---|---|---|---|---|
| Helicobacter pylori | 0.219 | Helicobacter pylori 26695-dR | 2 | 0.11 | 0.48 |
| Lactobacillus gasseri | 0.219 | Lactobacillus gasseri ATCC 33323 = JCM 1131 | 6 | 0.04 | 0.16 |
| Listeria monocytogenes | 0.219 | Listeria monocytogenes EGDe | 6 | 0.04 | 0.16 |
| Neisseria meningitidis | 0.219 | Neisseria meningitidis MC58 | 4 | 0.05 | 0.24 |
| Propionibacterium acnes | 0.219 | Cutibacterium acnes KPA171202 | 3 | 0.07 | 0.32 |
| Actinomyces odontolyticus | 0.02 | Actinomyces odontolyticus ATCC 17982 | 2 | 0.01 | 0.04 |
| Bacteroides vulgatus | 0.02 | Bacteroides vulgatus ATCC 8482 | 7 | 0.00 | 0.01 |
| Deinococcus radiodurans | 0.02 | Deinococcus radiodurans R1 | 3 | 0.01 | 0.03 |
| Enterococcus faecalis | 0.02 | Enterococcus faecalis OG1RF | 4 | 0.01 | 0.02 |
| Streptococcus pneumoniae | 0.02 | Streptococcus pneumoniae TIGR4 | 4 | 0.01 | 0.02 |

### Supplementary Note 1: RDP DB, Greengenes and SILVA statistics

The goal here is to count the number of unique genera, species and strains in all three databases for named high-quality named sequences. For RDP we use the training set for Naïve Bayes classifier. As the initial trusted sets (NOT representative set, but a set used for initial multiple sequence alignment) for Greengenes and SILVA are not available, we use Greengenes sequences clustered at 99% identity and SILVA “high-quality full length Ref dataset”. Then, we select only sequences that have species or strain annotations. Here we provide Linux shell commands to compare these databases.

#### RDP DB Training Set v16

For this part we will need to download both the training set and the full database (because the species / strain names are only present in the full database and that is the only piece of information used from it):

```
wget --no-check-certificate https://rdp.cme.msu.edu/download/current_Bacteria_unalign
ed.fa.gz
wget --no-check-certificate https://rdp.cme.msu.edu/download/current_Archaea_unaligne
d.fa.gz
gunzip *.gz
```

Download the original training set:

```
wget https://sourceforge.net/projects/rdp-classifier/files/RDP_Classifier_TrainingDat
a/RDPClassifier_16S_trainsetNo16_rawtrainingdata.zip/download
mv download rdp_trainset16.zip
unzip rdp_trainset16.zip
cd RDPClassifier_16S_trainsetNo16_rawtrainingdata
```

Extract RDP ID for each training set sequence, then use that to filter the entire RDP FASTA file:

```
grep "^>" trainset16_022016.fa | sed -E 's/.*\|([^\s]+)\sRoot.*$/\1/' > trainset16_rd
pids.txt
```

Get all strains:

```
grep "^>" current_Bacteria_unaligned.fa | cut -f1 | sed -E 's/^>([A-Z0-9]+) /\1\t/' |
sed -E 's/;//g' > rdp_v16_all_strains_bacteria.txt

grep "^>" current_Archaea_unaligned.fa | cut -f1 | sed -E 's/^>([A-Z0-9]+) /\1\t/' |
sed -E 's/;//g' > rdp_v16_all_strains_archaea.txt
```

```
cat rdp_v16_all_strains_bacteria.txt rdp_v16_all_strains_archaea.txt > rdp_v16_all_st
rains.txt
```

Now filter out strains with unknown names:

```
awk '$2 !~ "^[a-z]" && $2 !~ "^Bacterium" { print $0 }' rdp_v16_all_strains.txt > rdp
_v16_all_strains_with_names.txt

awk 'BEGIN {
  FS = OFS = "\t"
  }
NR == FNR {
  # while reading the 1st file
  # store its records in the array f
  f[$1] = $0
  next
  }
$1 in f {
  # when match is found
  # print all values
  print f[$1]
  }' rdp_v16_all_strains_with_names.txt \
  RDPClassifier_16S_trainsetNo16_rawtrainingdata/trainset16_rdpids.txt > \
  rdp_v16_train_strain_names.txt
```

Subtract 1 because the first line is blank to get number of unique strains

```
cut -f2 rdp_v16_train_strain_names.txt | sort | uniq > rdp_v16_trainset_unique_strain
s.txt
wc -l rdp_v16_trainset_unique_strains.txt
```

Unique species:

```
tail -n +2 rdp_v16_trainset_unique_strains.txt | cut -d" " -f1-2 | sort | uniq | wc -
l
```

Unique genera:

```
tail -n +2 rdp_v16_trainset_unique_strains.txt | cut -d" " -f1 | sort | uniq | wc -l
```

Unique: 12089 strains, 11675 species, 2465 genera

#### Greengenes 13.8

Download and extract:

```
wget ftp://greengenes.microbio.me/greengenes_release/gg_13_5/gg_13_8_otus.tar.gz
tar -xvzf gg_13_8_otus.tar.gz
```

Generate names with full species and count from sequences clustered at 99% identity:

```
grep -v "s__$" gg_13_8_otus/taxonomy/99_otu_taxonomy.txt | cut -d" " -f6,7 | sed -E '
s/([gs]__|\;)//g' | sort | uniq > gg_13_8_99otu_taxonomy_species.txt

sort gg_13_8_99otu_taxonomy_species.txt | uniq -c | wc -l
# 3114
```

```
# Now count unique genera (6th field, separated by space)
grep -v "g__;" gg_13_8_otus/taxonomy/99_otu_taxonomy.txt | cut -d " " -f6 | sed -E 's
/([gs]__|\;)//g' | grep "^[A-Z][a-z]" | sort | uniq -c | wc -l
# 1889
```

There is no strain information here. Uniques: NA strains, 3114 species, 1889 genera

#### SILVA v132

Generating statistics based on the SILVA_132_SSURef_tax_silva.fasta. First extract all entries that have species identification:

```
wget https://www.arb-silva.de/fileadmin/silva_databases/release_132/Exports/SILVA_132
_SSURef_tax_silva.fasta.gz
gunzip SILVA_132_SSURef_tax_silva.fasta.gz
mkdir silva
mv SILVA_132_SSURef_tax_silva.fasta.gz silva

awk '$0 ~ "^>" {
    split($0, a, ";");
    split(a[1], king, " ");
    if (a[7] !~ "[Uu][nidentified|ncultured]" && a[7] !~ " [Bb]acterium" &&
        a[7] ~ " " && a[7] !~ "^[a-z]" && (king[2] == "Bacteria" || king[2] == "Archa
ea") ) {
            sub("Candidatus[ ]?", "", a[7]);
            sub("\\[", "", a[7]);
            sub("\\]", "", a[7]);
            print a[7]
    }
}' SILVA_132_SSURef_tax_silva.fasta | sed -E s/\'//g | sort | uniq > SILVA_132_SSURe
f_tax_silva_species.txt
```

Also generate a single-line FASTA file with the same species:

```
awk ' {
    if ($0 ~ "^>") {
        split($0, a, ";");
        if (a[7] !~ "[Uu][nidentified|ncultured]" && a[7] !~ " [Bb]acterium" &&
            a[7] !~ "Bacterium" && a[7] !~ "^[Bb]acterium" && a[7] ~ " " &&
            a[7] !~ "^[a-z]") {
                sub("Candidatus[ ]?", "", a[7]);
                sub("\\[", "", a[7]);
                sub("\\]", "", a[7]);
                numrec += 1
                lastgood = 1
                if (numrec == 1) print ">"a[7];
                else print "\n>"a[7];
        } else {
            lastgood = 0;
        }
    } else {
        if (lastgood == 1) {
            gsub("U", "T", $0);
            printf $0;
        }
```

```
    }
}' SILVA_132_SSURef_tax_silva.fasta > SILVA_132_SSURef_tax_silva_species.fasta
```

This will give us a list of unique strains and species that have no strain designation. Extract only strains:

```
awk '$0 ~ "[^ ]+ [^ ]+ .*" { print $0 }' SILVA_132_SSURef_tax_silva_species.txt | sor
t | uniq > SILVA_132_SSURef_tax_silva_strains.txt
```

Also get a FASTA with strain names:

```
awk '{
    if ($0 ~ "^>") {
        n = split($0, a, " ");
        if (n > 2) {
            lastgood = 1;
            print $0;
        } else {
            lastgood = 0;
        }
    } else {
        if (lastgood == 1) print $0;
    }
}' SILVA_132_SSURef_tax_silva_species.fasta > SILVA_132_SSURef_tax_silva_strains.fast
a
```

Now generate a BLAST database from this FASTA file. We will use this database to match strain sequences from HiMAP database to this and look for 100% hits to see which strains have an exact match:

```
makeblastdb -dbtype nucl -in SILVA_132_SSURef_tax_silva_strains.fasta -out SILVA_132_
SSURef_tax_silva_strains
```

Count unique species assignments (replace multiple spaces from uniq with just one for easy manipulation):

```
cut -d" " -f1-2 SILVA_132_SSURef_tax_silva_species.txt | sed -E s/\'//g | grep -v "s
p\.$" | sort | uniq -c > SILVA_132_SSURef_tax_silva_species_unique_counts.txt

sed -E "s/[ ]+/ /g" SILVA_132_SSURef_tax_silva_species_unique_counts.txt > SILVA_132_
SSURef_tax_silva_species_unique_counts_table.txt
```

Count unique genera:

```
cut -d" " -f3 SILVA_132_SSURef_tax_silva_species_unique_counts_table.txt | sort | uni
q -c | sed -E 's/[ ]+/ /g' > SILVA_132_SSURef_tax_silva_genus_unique_counts_table.txt
```

Count unique strains:

```
sort SILVA_132_SSURef_tax_silva_strains.txt | uniq -c | wc -l
```

Uniques: 3242 genera, 12734 species, 116543 strains

Also generate a single-line FASTA file with the same species:

```
awk ' {
    if ($0 ~ "^>") {
        split($0, a, ";");
        if (a[7] !~ "[Uu][nidentified|ncultured]" && a[7] !~ " [Bb]acterium" &&
```

```
            a[7] !~ "Bacterium" && a[7] !~ "^[Bb]acterium" && a[7] ~ " " &&
            a[7] !~ "^[a-z]") {
                sub("Candidatus[ ]?", "", a[7]);
                sub("\\[", "", a[7]);
                sub("\\]", "", a[7]);
                numrec += 1
                lastgood = 1
                if (numrec == 1) print $0;
                else print "\n"$0;
        } else {
            lastgood = 0;
        }
    } else {
        if (lastgood == 1) {
            gsub("U", "T", $0);
            printf $0;
        }
    }
 }' SILVA_132_SSURef_tax_silva.fasta > SILVA_132_SSURef_tax_silva_species_wtax.fasta
```

What is the length distribution of these reference sequences? Let's use this for determining BLAST word size in one of the next steps.

```
awk '$0 !~ "^>" { print length($0) }' SILVA_132_SSURef_tax_silva_species_wtax.fasta |
sort -g | uniq -c | less
```

Most are between 1200 and 1500. Exact matching first between SILVA and GG was then used to find alignments between 100% hits. Generate a BLAST database:

```
makeblastdb -dbtype nucl -in SILVA_132_SSURef_tax_silva_species_wtax.fasta -out SILVA
_132_SSURef_tax_silva_species_wtax
```

Then BLAST the SILVA fasta file against this database.

#### Supplementary Note 2: PCR primers can reduce identity in 16S data

We analyzed the errors introduced by PCR primer sequences in the Zheng et al. 2015 mock dataset (from the main text). Even though a reverse PCR primer sequence is 5'-GGATTAGATACCC**BD**GTAGTC-3' (B = {C, G, T}, D = {A, G, T}), the most common sequence in the data is GGATTAGATACCC**CA**GTAGTC and not the correct GGATTAGATACCC**TG**GTAGTC for *Streptococcus mutans* (same problem occurs for every other species' sequence in this dataset):

```
Query  1    CCTACGGGAGGCAGCAGTAGGGAATCTTCGGCAATGGACGAAAGTCTGACCGAGCAACGC  60
            ||||||||||||||||||||||||||||||||||||||||||||||||||||||||||||
Sbjct  4    CCTACGGGAGGCAGCAGTAGGGAATCTTCGGCAATGGACGAAAGTCTGACCGAGCAACGC  63

Query  61   CGCGTGAGTGAAGAAGGTTTTCGGATCGTAAAGCTCTGTTGTAAGTCAAGAACGTGTGTG  120
            ||||||||||||||||||||||||||||||||||||||||||||||||||||||||||||
Sbjct  64   CGCGTGAGTGAAGAAGGTTTTCGGATCGTAAAGCTCTGTTGTAAGTCAAGAACGTGTGTG  123

Query  121  AGAGTGGAAAGTTCACACAGTGACGGTAGCTTACCAGAAAGGGACGGCTAACTACGTGCC  180
            ||||||||||||||||||||||||||||||||||||||||||||||||||||||||||||
```

```
Sbjct  124  AGAGTGGAAAGTTCACACAGTGACGGTAGCTTACCAGAAAGGGACGGCTAACTACGTGCC  183

Query  181  AGCAGCCGCGGTAATACGTAGGTCCCGAGCGTTGTCCGGATTTATTGGGCGTAAAGGGAG  240
            ||||||||||||||||||||||||||||||||||||||||||||||||||||||||||||
Sbjct  184  AGCAGCCGCGGTAATACGTAGGTCCCGAGCGTTGTCCGGATTTATTGGGCGTAAAGGGAG  243

Query  241  CGCAGGCGGTCAGGAAAGTCTGGAGTAAAAGGCTATGGCTCAACCATAGTGTGCTCTGGA  300
            ||||||||||||||||||||||||||||||||||||||||||||||||||||||||||||
Sbjct  244  CGCAGGCGGTCAGGAAAGTCTGGAGTAAAAGGCTATGGCTCAACCATAGTGTGCTCTGGA  303

Query  301  AACTGTCTGACTTGAGTGCAGAAGGGGAGAGTGGAATTCCATGTGTAGCGGTGAAATGCG  360
            ||||||||||||||||||||||||||||||||||||||||||||||||||||||||||||
Sbjct  304  AACTGTCTGACTTGAGTGCAGAAGGGGAGAGTGGAATTCCATGTGTAGCGGTGAAATGCG  363

Query  361  TAGATATATGGAGGAACACCAGTGGCGAAAGCGGCTCTCTGGTCTGTCACTGACGCTGAG  420
            ||||||||||||||||||||||||||||||||||||||||||||||||||||||||||||
Sbjct  364  TAGATATATGGAGGAACACCAGTGGCGAAAGCGGCTCTCTGGTCTGTCACTGACGCTGAG  423

Query  421  GCTCGAAAGCGTGGGTAGCGAACAGGATTAGATACCCCAGTAGTC  465
            |||||||||||||||||||||||||||||||||||||  ||||||
Sbjct  424  GCTCGAAAGCGTGGGTAGCGAACAGGATTAGATACCCTGGTAGTC  468
```

Top sequence from the data (Query), bottom is the reference exact sequence (Subject). Forward PCR primer region is shown in green and the reverse PCR primer region is shown in red. This may be caused by an uneven concentration of different primer sequences.

#### Supplementary Note 3: DADA2, QIIME2 and QIIME1 pipelines

Following workflow recommendations for each pipeline, DADA2 was used with SILVA v132 NR database, QIIME2 with Greengenes database clustered at 99% identity and Deblur denoiser, and QIIME1 with Greengenes database clustered at 97% identity both using Open Reference and De Novo OTU picking. The main text presents the (recommended) workflow using Open Reference OTUs.

##### Mock: DADA2 SILVA pipeline

```
#--------------- DADA2 pipeline for Zheng 2015 et al. --------------------
suppressPackageStartupMessages(library(data.table))
suppressPackageStartupMessages(library(dada2))

main_path = '~/data1/'
set_path = file.path(main_path, 'zheng-2015')
setwd(set_path)

# Follow the DADA2 pipeline:
# https://benjjneb.github.io/dada2/tutorial.html

# Define some helper functions
fasta_writer = function (meta, seqs, output, ncpu=detectCores()-1) {
  # Write FASTA sequences from meta data and sequences
```

```
  writeChar(paste(paste0('>', meta),
                  seqs,
                  sep='\n', collapse='\n'
            ),
            output, eos=NULL)
}

#--------------- Filter and trim files ----------------------------------
fq_path = file.path(set_path,'fastq_v3v4')
fq_fwd = sort(list.files(fq_path, pattern='.*R1.*gz', full.names=T))
fq_rev = sort(list.files(fq_path, pattern='.*R2.*gz', full.names=T))
sample_ids = sapply(strsplit(basename(fq_fwd), '_', fixed=T), `[`, 1)
# Check quality profiles
plotQualityProfile(fq_fwd[1:2])
plotQualityProfile(fq_rev[1:2])
# Trim last 100 nts for reverse reads. Generate output filenames:
fq_fwd_fil = file.path(set_path, 'dada2_analysis/filtered',
                       paste0(sample_ids, '_R1_filtered.fastq'))
fq_rev_fil = file.path(set_path, 'dada2_analysis/filtered',
                       paste0(sample_ids, '_R2_filtered.fastq'))
# Filter and trim
ft_out = filterAndTrim(fq_fwd, fq_fwd_fil, fq_rev, fq_rev_fil,
                       trimLeft=c(22,22), truncLen=c(300,200), maxN=0,
maxEE=c(2,2),
                       truncQ=2, rm.phix=T, compress=T, multithread=T)
# Check the number of retained reads after paired filter
head(ft_out)
# Most reads retained.

#--------------------- DADA2 denoising ----------------------------------
# Learn errors for fwd and rev reads separately
# This step takes few hours on 2016 Macbook Pro. In my tests with other
# data sets, it works equally well to use much less than 1e6 reads for
# learning error rates. 10-100k read sample was still fine.
err_fwd = learnErrors(fq_fwd_fil, multithread=T)
err_rev = learnErrors(fq_rev_fil, multithread=T)

# Save intermediate R files, so we can resume later if needed.
saveRDS(err_fwd, 'dada2_analysis/err_fwd')
saveRDS(err_rev, 'dada2_analysis/err_rev')

# Plot errors to check how well learned errors fit
plotErrors(err_fwd, nominalQ=T)
plotErrors(err_rev, nominalQ=T)

# Looks good. Run dada2.
derepFs = derepFastq(fq_fwd_fil, verbose=T)
derepRs = derepFastq(fq_rev_fil, verbose=T)
# Name the derep-class objects by the sample names
names(derepFs) = sample_ids
```

```
names(derepRs) = sample_ids
dadaFs = dada(derepFs, err=err_fwd, multithread=T)
dadaRs = dada(derepRs, err=err_rev, multithread=T)
saveRDS(dadaFs, 'dada2_analysis/dadaFs')
saveRDS(dadaRs, 'dada2_analysis/dadaRs')

# Merge reads
mergers = mergePairs(dadaFs, derepFs, dadaRs, derepRs, verbose=T)
saveRDS(mergers, 'dada2_analysis/mergers')

# Generate table with sequence counts
seqtab = makeSequenceTable(mergers)
saveRDS(seqtab, 'dada2_analysis/seqtab')

# Chimera removal
seqtab.nochim = removeBimeraDenovo(seqtab, method="consensus",
                                   multithread=T, verbose=T)
saveRDS(seqtab.nochim, 'dada2_analysis/seqtab.nochim')
# For multiple samples collapse:
# seqtab.nochim.coll = collapseNoMismatch(seqtab.nochim)

# Fraction of chimeric reads
cat('Fraction of non-chimeric reads: ', sum(seqtab.nochim)/sum(seqtab), '\n')
# 94%. That looks good.

# Assign taxonomy with SILVA
taxa = assignTaxonomy(seqtab.nochim,
                      'dada2_analysis/db/silva_nr_v132_train_set.fa.gz',
                      multithread=T, verbose=T)
saveRDS(taxa, 'dada2_analysis/taxa_before_addSpecies')
# Add species assignment
taxa = addSpecies(taxa,
'dada2_analysis/db/silva_species_assignment_v132.fa.gz',
                  allowMultiple=T, verbose=T)
saveRDS(taxa, 'dada2_analysis/taxa')

# Print assignments
taxa.print = taxa
rownames(taxa.print) = NULL

# Save taxonomy abundance and FASTA sequences
tax.dt = as.data.table(taxa, keep.rownames=T)
write.table(tax.dt, 'dada2_analysis/taxonomy.txt',
            sep='\t', row.names=F, quote=F)
ab.dt = as.data.table(t(seqtab.nochim), keep.rownames=T)
write.table(ab.dt,
            'dada2_analysis/abundances.txt',
            sep='\t', row.names=F, quote=F)

# Write FASTA file with sequences
seqs.dt = ab.dt[, .(rn, id=1:.N)]
```

```
fasta_writer(seqs.dt[, id], seqs.dt[, rn],
              'dada2_analysis/sequences.fasta')

# We manually BLAST these 20 sequences against NCBI database to identify
# the 20 mock species/strains.
# blastn -query sequences.fasta \
#        -db /data1/igor/himap/db/V3-V4_341F-805R_hang22_uniq \
#        -num_threads 4 -outfmt 6 > sequences_blast.txt
# Then we used these assignments to check which sequence was found or missing.
```

#### Mock: QIIME2 GG99 pipeline

In the Zheng et al. 2015 mock community dataset PCR primers were still present (unlike the DIABIMMUNE data, where this step is skipped!). We found that trimming first 22 nt instead of using cutadapt (can used through QIIME2) in this case removes PCR primers more accurately. This is an important step, because we noticed that a PCR primer with a mismatch to the exact sequence can be more common than the one with an exact match. This single nt mismatch might make taxonomy prediction more difficult later down the pipeline.

```
# Trim PCR primers manually (tried cutadapt within QIIME2 and it doesn't remove all
of them for some reason)

mkdir fastq_trimmed
vsearch --fastx_filter ../fastq_v3v4/V3V4Rep1_S5_L001_R1_001.fastq.gz --
fastq_stripleft 22 --fastqout fastq_trimmed/V3V4Rep1_S5_L001_R1_001.fastq
vsearch --fastx_filter ../fastq_v3v4/V3V4Rep1_S5_L001_R2_001.fastq.gz --
fastq_stripleft 22 --fastqout fastq_trimmed/V3V4Rep1_S5_L001_R2_001.fastq
vsearch --fastx_filter ../fastq_v3v4/V3V4Rep2_S6_L001_R1_001.fastq.gz --
fastq_stripleft 22 --fastqout fastq_trimmed/V3V4Rep2_S6_L001_R1_001.fastq
vsearch --fastx_filter ../fastq_v3v4/V3V4Rep2_S6_L001_R2_001.fastq.gz --
fastq_stripleft 22 --fastqout fastq_trimmed/V3V4Rep2_S6_L001_R2_001.fastq
vsearch --fastx_filter ../fastq_v3v4/V3V4Rep3_S7_L001_R1_001.fastq.gz --
fastq_stripleft 22 --fastqout fastq_trimmed/V3V4Rep3_S7_L001_R1_001.fastq
vsearch --fastx_filter ../fastq_v3v4/V3V4Rep3_S7_L001_R2_001.fastq.gz --
fastq_stripleft 22 --fastqout fastq_trimmed/V3V4Rep3_S7_L001_R2_001.fastq
vsearch --fastx_filter ../fastq_v3v4/V3V4Rep4_S8_L001_R1_001.fastq.gz --
fastq_stripleft 22 --fastqout fastq_trimmed/V3V4Rep4_S7_L001_R1_001.fastq
vsearch --fastx_filter ../fastq_v3v4/V3V4Rep4_S8_L001_R2_001.fastq.gz --
fastq_stripleft 22 --fastqout fastq_trimmed/V3V4Rep4_S7_L001_R2_001.fastq

cd fastq_trimmed
gzip *.fastq
cd ..
```

Manually create a manifest.txt files:

```
sample-id,absolute-filepath,direction
V3V4Rep1,/Users/igor/cloud/research/microbiome/zheng-2015/qiime2_analysis/fastq_trimm
ed/V3V4Rep1_S5_L001_R1_001.fastq.gz,forward
V3V4Rep1,/Users/igor/cloud/research/microbiome/zheng-2015/qiime2_analysis/fastq_trimm
ed/V3V4Rep1_S5_L001_R2_001.fastq.gz,reverse
V3V4Rep2,/Users/igor/cloud/research/microbiome/zheng-2015/qiime2_analysis/fastq_trimm
ed/V3V4Rep2_S6_L001_R1_001.fastq.gz,forward
```

```
V3V4Rep2,/Users/igor/cloud/research/microbiome/zheng-2015/qiime2_analysis/fastq_trimm
ed/V3V4Rep2_S6_L001_R2_001.fastq.gz,reverse
V3V4Rep3,/Users/igor/cloud/research/microbiome/zheng-2015/qiime2_analysis/fastq_trimm
ed/V3V4Rep3_S7_L001_R1_001.fastq.gz,forward
V3V4Rep3,/Users/igor/cloud/research/microbiome/zheng-2015/qiime2_analysis/fastq_trimm
ed/V3V4Rep3_S7_L001_R2_001.fastq.gz,reverse
V3V4Rep4,/Users/igor/cloud/research/microbiome/zheng-2015/qiime2_analysis/fastq_trimm
ed/V3V4Rep4_S8_L001_R1_001.fastq.gz,forward
V3V4Rep4,/Users/igor/cloud/research/microbiome/zheng-2015/qiime2_analysis/fastq_trimm
ed/V3V4Rep4_S8_L001_R2_001.fastq.gz,reverse
```

```
# Import paired-end FASTQ files into artifact
qiime tools import \
    --type 'SampleData[PairedEndSequencesWithQuality]' \
    --input-path manifest_trimmed.txt \
    --output-path qiime_import_trimmed \
    --source-format PairedEndFastqManifestPhred33

# Merge reads
qiime vsearch join-pairs \
    --i-demultiplexed-seqs qiime_import_trimmed.qza \
    --o-joined-sequences qiime_import_trimmed_merged.qza

# Quality filter merged reads
qiime quality-filter q-score-joined \
    --i-demux qiime_import_trimmed_merged.qza \
    --o-filtered-sequences qiime_import_trimmed_merged_filtered.qza \
    --o-filter-stats qiime_import_trimmed_merged_filtered_stats.qza

# Run Deblur denoising
qiime deblur denoise-16S \
    --i-demultiplexed-seqs qiime_import_trimmed_merged_filtered.qza \
    --p-trim-length 393 \
    --o-representative-sequences qiime_import_trimmed_merged_filtered_rep-seqs.qza \
    --o-table qiime_import_trimmed_merged_filtered_table.qza \
    --p-sample-stats \
    --o-stats qiime_import_trimmed_merged_filtered_deblur_stats.qza

# Export data from artifacts into normal files
qiime tools export qiime_import_trimmed_merged_filtered_table.qza \
    --output-dir qiime_import_trimmed_merged_filtered_feature_table
qiime tools export qiime_import_trimmed_merged_filtered_rep-seqs.qza \
    --output-dir qiime_import_trimmed_merged_filtered_feature_table
biom convert -i qiime_import_trimmed_merged_filtered_feature_table/feature-table.biom
\
    -o qiime_import_trimmed_merged_filtered_feature_table/feature-table.txt --to-tsv
```

For taxonomic classification, we have to do a bit more work since we couldn't find precomputed artifact file for v3-v4 region on QIIME2 website. There's only precomputed files for either full-length 16S sequences or for weirdly short v4 region (120 nt?) at https://docs.qiime2.org/2018.2/data-resources/ (scroll a bit down).

Let’s follow the guide from here https://docs.qiime2.org/2018.2/tutorials/feature-classifier/ to train Naive Bayes classifier on V3-V4 region of 99% OTU Greengenes 13.8 sequences. First download Greengenes 13.8 99% OTU files:

```
# Download fasta and taxonomy from Greengenes
mkdir greengenes
cd greengenes
wget ftp://greengenes.microbio.me/greengenes_release/gg_13_5/gg_13_8_otus.tar.gz
tar xvfz gg_13_8_otus.tar.gz
rm gg_13_8_otus.tar.gz
mv gg_13_8_otus/rep_set/99_otus.fasta .
mv gg_13_8_otus/taxonomy/99_otu_taxonomy.txt .
rm -rf gg_13_8_otus
cd ..

# Import FASTA and txt files as QIIME2 artifacts
qiime tools import \
  --type 'FeatureData[Sequence]' \
  --input-path greengenes/99_otus.fasta \
  --output-path 99_otus.qza

qiime tools import \
  --type 'FeatureData[Taxonomy]' \
  --source-format HeaderlessTSVTaxonomyFormat \
  --input-path greengenes/99_otu_taxonomy.txt \
  --output-path ref-taxonomy.qza
```

We will also need primer sequences for V3-V4 region. Forward primer 341F 5’-3’ sequence: GACAGCCTACGGGNGGCWGCAG. Reverse primer 805R 3’-5’ sequence: GACTACCAGGGTATCTAATC. Now we can extract reference reads and truncate to 419 nt:

```
# Extract V3-V4 regions
qiime feature-classifier extract-reads \
    --i-sequences 99_otus.qza \
    --p-f-primer GACAGCCTACGGGNGGCWGCAG \
    --p-r-primer GACTACCAGGGTATCTAATC \
    --p-trunc-len 419 \
    --o-reads 99_otus_refseqs.qza

# Train the classifier
qiime feature-classifier fit-classifier-naive-bayes \
    --i-reference-reads 99_otus_refseqs.qza \
    --i-reference-taxonomy ref-taxonomy.qza \
    --o-classifier 99_otus_v3-v4_341f-805r_classifier.qza
```

Now run this feature classifier to generate taxonomy for the sequences in our data:

```
qiime feature-classifier classify-sklearn \
  --i-classifier 99_otus_v3-v4_341f-805r_classifier.qza \
  --i-reads qiime_import_trimmed_merged_filtered_rep-seqs.qza \
  --o-classification qiime_import_trimmed_merged_filtered_taxonomy.qza

# Export taxonomy to tab-delimited file
qiime tools export qiime_import_trimmed_merged_filtered_taxonomy.qza \
  --output-dir qiime_import_trimmed_merged_filtered_taxonomy
```

#### Mock: QIIME1 GG97 pipeline

Start macOS session using `macqiime`. This uses QIIME 1.9. Run everything from zheng-2015 folder. USEARCH 6.1 needs to be manually downloaded from http://www.drive5.com and be present in PATH as `usearch61` executable for the chimera removal part.

##### Merge reads

Start from PCR primer trimmed reads that we prepared from QIIME 2 analysis. The merging is done using (the default) `fastq-join` algorithm:

```
join_paired_ends.py -f qiime2_analysis/fastq_trimmed/V3V4Rep1_S5_L001_R1_001.fastq.gz
\
                    -r qiime2_analysis/fastq_trimmed/V3V4Rep1_S5_L001_R2_001.fastq.gz
\
                    -o qiime1_analysis/trimmed_merged/V3V4Rep1_merged.fastq

join_paired_ends.py -f qiime2_analysis/fastq_trimmed/V3V4Rep2_S6_L001_R1_001.fastq.gz
\
                    -r qiime2_analysis/fastq_trimmed/V3V4Rep2_S6_L001_R2_001.fastq.gz
\
                    -o qiime1_analysis/trimmed_merged/V3V4Rep2_merged.fastq

join_paired_ends.py -f qiime2_analysis/fastq_trimmed/V3V4Rep3_S7_L001_R1_001.fastq.gz
\
                    -r qiime2_analysis/fastq_trimmed/V3V4Rep3_S7_L001_R2_001.fastq.gz
\
                    -o qiime1_analysis/trimmed_merged/V3V4Rep3_merged.fastq

join_paired_ends.py -f qiime2_analysis/fastq_trimmed/V3V4Rep4_S8_L001_R1_001.fastq.gz
\
                    -r qiime2_analysis/fastq_trimmed/V3V4Rep4_S8_L001_R2_001.fastq.gz
\
                    -o qiime1_analysis/trimmed_merged/V3V4Rep4_merged.fastq
```

##### Quality control and filtering

```
split_libraries_fastq.py -i qiime1_analysis/trimmed_merged/V3V4Rep1_merged.fastq/fast
qjoin.join.fastq \
                         --sample_ids V3V4Rep1 \
                         -o qiime1_analysis/V3V4Rep1_quality_filtered_q20/ -q 19 \
                         --barcode_type 'not-barcoded' --phred_offset=33

split_libraries_fastq.py -i qiime1_analysis/trimmed_merged/V3V4Rep2_merged.fastq/fast
qjoin.join.fastq \
                         --sample_ids V3V4Rep2 \
                         -o qiime1_analysis/V3V4Rep2_quality_filtered_q20/ -q 19 \
                         --barcode_type 'not-barcoded' --phred_offset=33

split_libraries_fastq.py -i qiime1_analysis/trimmed_merged/V3V4Rep3_merged.fastq/fast
qjoin.join.fastq \
                         --sample_ids V3V4Rep3 \
                         -o qiime1_analysis/V3V4Rep3_quality_filtered_q20/ -q 19 \
                         --barcode_type 'not-barcoded' --phred_offset=33
```

```
split_libraries_fastq.py -i qiime1_analysis/trimmed_merged/V3V4Rep4_merged.fastq/fast
qjoin.join.fastq \
                         --sample_ids V3V4Rep4 \
                         -o qiime1_analysis/V3V4Rep4_quality_filtered_q20/ -q 19 \
                         --barcode_type 'not-barcoded' --phred_offset=33
```

#### Chimera removal

```
wget https://drive5.com/uchime/gold.fa --no-check-certificate

# Make sure usearch61 (this needs to be the name of the executable) is in %PATH%
identify_chimeric_seqs.py -m usearch61 -i V3V4Rep1_quality_filtered_q20/seqs.fna -r g
old.fa -o qiime_chimeras1/
identify_chimeric_seqs.py -m usearch61 -i V3V4Rep2_quality_filtered_q20/seqs.fna -r g
old.fa -o qiime_chimeras2/
identify_chimeric_seqs.py -m usearch61 -i V3V4Rep3_quality_filtered_q20/seqs.fna -r g
old.fa -o qiime_chimeras3/
identify_chimeric_seqs.py -m usearch61 -i V3V4Rep4_quality_filtered_q20/seqs.fna -r g
old.fa -o qiime_chimeras4/

mkdir qiime_nochim
filter_fasta.py -f V3V4Rep1_quality_filtered_q20/seqs.fna \
                -o qiime_nochim/V3V4Rep1_nochim.fasta \
                -s qiime_chimeras1/chimeras.txt -n

filter_fasta.py -f V3V4Rep2_quality_filtered_q20/seqs.fna \
                -o qiime_nochim/V3V4Rep2_nochim.fasta \
                -s qiime_chimeras2/chimeras.txt -n

filter_fasta.py -f V3V4Rep3_quality_filtered_q20/seqs.fna \
                -o qiime_nochim/V3V4Rep3_nochim.fasta \
                -s qiime_chimeras3/chimeras.txt -n

filter_fasta.py -f V3V4Rep4_quality_filtered_q20/seqs.fna \
                -o qiime_nochim/V3V4Rep4_nochim.fasta \
                -s qiime_chimeras4/chimeras.txt -n
```

#### Pick open reference OTUs

```
pick_open_reference_otus.py -i qiime_nochim/V3V4Rep1_nochim.fasta -o V3V4Rep1_openref
_otus
pick_open_reference_otus.py -i qiime_nochim/V3V4Rep2_nochim.fasta -o V3V4Rep2_openref
_otus
pick_open_reference_otus.py -i qiime_nochim/V3V4Rep3_nochim.fasta -o V3V4Rep3_openref
_otus
pick_open_reference_otus.py -i qiime_nochim/V3V4Rep4_nochim.fasta -o V3V4Rep4_openref
_otus
```

#### Pick de novo OTUs

We show Open reference OTU picking in the main text, but de novo OTU picking results look very similar.

```
pick_de_novo_otus.py -i qiime_nochim/V3V4Rep1_nochim.fasta -o V3V4Rep1_denovo_otus
pick_de_novo_otus.py -i qiime_nochim/V3V4Rep2_nochim.fasta -o V3V4Rep2_denovo_otus
```

```
pick_de_novo_otus.py -i qiime_nochim/V3V4Rep3_nochim.fasta -o V3V4Rep3_denovo_otus
pick_de_novo_otus.py -i qiime_nochim/V3V4Rep4_nochim.fasta -o V3V4Rep4_denovo_otus
```

##### Extract counts for each sample id

For open ref OTUs:

```
awk '{ print $1, NF-1 }' V3V4Rep1_openref_otus/final_otu_map_mc2.txt > V3V4Rep1_otu_c
ounts_openref.txt
awk '{ print $1, NF-1 }' V3V4Rep2_openref_otus/final_otu_map_mc2.txt > V3V4Rep2_otu_c
ounts_openref.txt
awk '{ print $1, NF-1 }' V3V4Rep3_openref_otus/final_otu_map_mc2.txt > V3V4Rep3_otu_c
ounts_openref.txt
awk '{ print $1, NF-1 }' V3V4Rep4_openref_otus/final_otu_map_mc2.txt > V3V4Rep4_otu_c
ounts_openref.txt
```

For de novo OTUs:

```
awk '{ print $1, NF-1 }' V3V4Rep1_denovo_otus/uclust_picked_otus/V3V4Rep1_nochim_otus
.txt > V3V4Rep1_denovo_otus/otu_counts.txt
awk '{ print $1, NF-1 }' V3V4Rep2_denovo_otus/uclust_picked_otus/V3V4Rep2_nochim_otus
.txt > V3V4Rep2_denovo_otus/otu_counts.txt
awk '{ print $1, NF-1 }' V3V4Rep3_denovo_otus/uclust_picked_otus/V3V4Rep3_nochim_otus
.txt > V3V4Rep3_denovo_otus/otu_counts.txt
awk '{ print $1, NF-1 }' V3V4Rep4_denovo_otus/uclust_picked_otus/V3V4Rep4_nochim_otus
.txt > V3V4Rep4_denovo_otus/otu_counts.txt
```

#### DIABIMMUNE: DADA2 SILVA pipeline

Used latest (at the time of writing) DADA2 v1.8, analysis tutorial followed at https://benjjneb.github.io/dada2/tutorial.html (accessed May 14th, 2018). As per instructions, we downloaded *silva_nr_v132_train_set.fa.gz* and *silva_species_assignment_v132.fa.gz* files for taxonomic classification. The following is the R code used to produce results in the main text:

```
library(dada2)
library(data.table)

main_path = '/data1/igor/diabimmune'
fq_fwd = sort(dir(file.path(main_path, '16s_fastq_all'), 'R1', full.names=T))
fq_rev = sort(dir(file.path(main_path, '16s_fastq_all'), 'R2', full.names=T))

sample.names = sapply(strsplit(basename(fq_fwd), "_"), `[`, 1)

# Use this to guesstimte truncLen
plotQualityProfile(fq_fwd[1:2])
plotQualityProfile(fq_rev[1:2])

# Filter and trim
filtFs = file.path(main_path, 'dada2', 'filtered', paste0(sample.names,
"_F_filt.fastq.gz"))
filtRs = file.path(main_path, 'dada2', 'filtered', paste0(sample.names,
"_R_filt.fastq.gz"))
filtOut = filterAndTrim(fq_fwd, filtFs, fq_rev, filtRs, truncLen=c(150,150),
```

```
                         maxN=0, maxEE=c(2,2), truncQ=2, rm.phix=TRUE,
                         compress=TRUE, multithread=TRUE)

# Learn errors
errF = learnErrors(filtFs, multithread=TRUE)
errR = learnErrors(filtRs, multithread=TRUE)

# Iterate 1 by 1 sample since there are too many to load all into memory
dadaFs = list()
dadaRs = list()
mergers = list()
for (i in 1:length(sample.names)) {
  # Forward read
  cat('Processing sample ', i , ' out of ', length(sample.names), '\n')
  derepFs = list(derepFastq(filtFs[i], verbose=F))
  names(derepFs) = sample.names[i]
  dadaFs[[i]] = dada(derepFs[[1]], err=errF, multithread=TRUE)
  # Reverse read
  derepRs = list(derepFastq(filtRs[i], verbose=F))
  names(derepRs) = sample.names[i]
  dadaRs[[i]] = dada(derepRs[[1]], err=errR, multithread=TRUE)
  # Merge
  mergers[[i]] = mergePairs(dadaFs[[i]], derepFs[[1]], dadaRs[[i]],
derepRs[[1]], verbose=TRUE)
}

# Save DADA results
saveRDS(dadaFs, file.path(main_path, 'dada2', 'dadaFs'))
saveRDS(dadaRs, file.path(main_path, 'dada2', 'dadaRs'))
saveRDS(mergers, file.path(main_path, 'dada2', 'mergers'))

# Generate sequence count table
seqtab = makeSequenceTable(mergers)
saveRDS(seqtab, file.path(main_path, 'dada2', 'seqtab'))
dim(seqtab)

# Remove chimeras
seqtab.nochim = removeBimeraDenovo(seqtab, method="consensus",
multithread=TRUE, verbose=TRUE)
rownames(seqtab.nochim) = sample.names
saveRDS(seqtab.nochim, file.path(main_path, 'dada2', 'seqtab.nochim'))

# Assign taxonomy
taxa = assignTaxonomy(seqtab.nochim,
'/data1/igor/databases/silva_nr_v132_train_set.fa.gz',
                      multithread=TRUE)
taxa = addSpecies(taxa,
'/data1/igor/databases/silva_species_assignment_v132.fa.gz',
                  allowMultiple=T)
saveRDS(taxa, file.path(main_path, 'dada2', 'taxa'))
```

```
# Convert matrices to data tables
seqtab.dt = melt(as.data.table(seqtab.nochim, keep.rownames=T),
                 variable.name='sequence', value.name='count', id.vars='rn')
seqtab.dt[, seq_id := .GRP, by=sequence]
seqtab.dt = seqtab.dt[count > 0]
seqtab.dt[, sequence := NULL]

taxa.dt = as.data.table(taxa, keep.rownames=T)
taxa.dt = merge(taxa.dt, unique(seqtab.dt[, .(seq_id, sequence)]),
                by.x='rn', by.y='sequence')
taxa.dt[, rn := NULL]
setorderv(taxa.dt, c('seq_id'))

# Write tables
write.table(seqtab.dt, file.path(main_path, 'dada2', 'seqtab.dt.txt'),
            sep='\t', quote=F, row.names=F)

write.table(taxa.dt, file.path(main_path, 'dada2', 'taxa.dt.txt'),
            sep='\t', quote=F, row.names=F)
```

#### DIABIMMUNE: QIIME2 GG99 pipeline

##### Manifest

The analysis was done using QIIME version 2018.4. First we need to generate a manifest file (metadata) for QIIME2 to use. We generate it from the raw FASTQ files (from the folder 16_fastq_all) using this bash script “generate_manifest”:

```
#!/bin/bash
#
# Arguments: $1 absolute filepath to scan for fastq files, e.g.
"/data1/igor/diabimmune/16s_fastq_all/"
#            $2 separator for sample id, e.g. "_"
#            $3 forward read substring, e.g. "_R1"
#            $4 reverse reads substring, e.g. "_R2"

echo "sample-id,absolute-filepath,direction"
for f in $(find $1 -name "*.fastq*"); do
        base=${f##*/}
        if [[ $base = *"$3"* ]]; then
                direction="forward"
        fi
        if [[ $base = *"$4"* ]]; then
                direction="reverse"
        fi
        echo "${base%%_*},$f,$direction"

done
```

Now run the script:

```
./generate_manifest /data1/igor/diabimmune/16s_fastq_all/ _ _R1 _R2 > manifest.txt
```

Import FASTQ files into artifact:

```
# This part takes A LONG time (about a day and a half on our server)
qiime tools import \
    --type 'SampleData[PairedEndSequencesWithQuality]' \
    --input-path manifest.txt \
    --output-path diab_import \
    --source-format PairedEndFastqManifestPhred33

# This is a large dataset and QIIME2 copies stuff over and over into and out
# of the temp folder, which will run out of space on our 50GB boot partition.
# Change the temp folder to the large storage volume.
export TMPDIR="/data1/igor/diabimmune/qiime2/tmp/"

# Merge reads
qiime vsearch join-pairs \
    --i-demultiplexed-seqs diab_import.qza \
    --o-joined-sequences diab_merged.qza

# Quality filter merged reads
qiime quality-filter q-score-joined \
    --i-demux diab_merged.qza \
    --o-filtered-sequences diab_filtered.qza \
    --o-filter-stats diab_filtered_stats.qza

# Run Deblur denoising
qiime deblur denoise-16S \
    --i-demultiplexed-seqs diab_filtered.qza \
    --p-trim-length 251 \
    --o-representative-sequences diab_repseqs.qza \
    --o-table diab_table.qza \
    --p-sample-stats \
    --o-stats diab_deblur_stats.qza

# Export data from artifacts into normal files
qiime tools export diab_table.qza \
    --output-dir export_feature_table
qiime tools export diab_repseqs.qza \
    --output-dir export_feature_table
biom convert -i export_feature_table/feature-table.biom \
        -o diab_feature-table.txt --to-tsv
```

For taxonomic classification, we have to do a bit more work since I couldn't find precomputed artifact file for the normal V4 region on QIIME2 website. There's only precomputed files for either full-length 16S sequences or for unusually short v4 region (120 nt) at https://docs.qiime2.org/2018.2/data-resources/ (scroll a bit down).

Let's follow the guide from here https://docs.qiime2.org/2018.2/tutorials/feature-classifier/ to train Naive Bayes classifier on V4 region of 99% OTU Greengenes 13.8 sequences. Download the Greengenes 13.8 99% OTU files (FASTA from rep_set and taxonomy) into greengenes folder and then run this:

```
# Import FASTA and txt files as QIIME2 artifacts
qiime tools import \
  --type 'FeatureData[Sequence]' \
  --input-path greengenes/99_otus.fasta \
  --output-path 99_otus.qza
```

```
qiime tools import \
  --type 'FeatureData[Taxonomy]' \
  --source-format HeaderlessTSVTaxonomyFormat \
  --input-path greengenes/99_otu_taxonomy.txt \
  --output-path ref-taxonomy.qza
```

We will also need primer sequences for V4 region. Forward primer 515F 5'-3' sequence: GTGCCAGCMGCCGCGGTAA. Reverse primer 805R 3'-5' sequence: GACTACCAGGGTATCTAATC. Now we can extract reference reads and truncate to 251 nt because that's what we used in the HiMAP pipeline:

```
# Extract V3-V4 regions
qiime feature-classifier extract-reads \
    --i-sequences 99_otus.qza \
    --p-f-primer GTGCCAGCMGCCGCGGTAA \
    --p-r-primer GACTACCAGGGTATCTAATC \
    --p-trunc-len 251 \
    --o-reads 99_otus_refseqs.qza

# Train the classifier
qiime feature-classifier fit-classifier-naive-bayes \
    --i-reference-reads 99_otus_refseqs.qza \
    --i-reference-taxonomy ref-taxonomy.qza \
    --o-classifier 99_otus_v4_515f-805r_classifier.qza
```

Now run this feature classifier to generate taxonomy for the sequences in our data:

```
qiime feature-classifier classify-sklearn \
  --i-classifier 99_otus_v4_515f-805r_classifier.qza \
  --i-reads diab_repseqs.qza \
  --o-classification diab_taxonomy.qza

# Export taxonomy to tab-delimited file
qiime tools export diab_taxonomy.qza \
  --output-dir export_taxonomy
```

#### DIABIMMUNE: QIIME1 GG97 pipeline

All raw paired-end FASTQ files (*R1* and *R2*) are placed in "16s_fastq_all" folder. Then the following commands were ran. The analysis was done using QIIME version 1.9 (as part of the Anaconda installation), ran natively under Ubuntu Linux 16.04.4 LTS.

##### Merge reads

```
mkdir merged
# This will generate a separate folder for each read pair
for f in $(find ../16s_fastq_all -name "*_R1*.fastq*"); do
    base=${f##*/}
    base=${base%%_*}
    join_paired_ends.py -f $f \
                    -r ${f/_R1/_R2} \
                    -o merged/${base}
done
```

```
# Extract files from all folders into merged, cleanup unmerged
for f in $(find merged -name "*.join.fastq"); do
    base=${f#*/}
    base=${base%%/*}
    mv $f merged/${base}.fastq
    rm -rf merged/${base}
done
```

Quality filtering

```
mkdir filtered_q20
for f in $(find merged -maxdepth 1 -name "*.fastq"); do
    base=${f%.*} # remove extension
    base=${base#*/}
    echo $f
    echo $base
    split_libraries_fastq.py -i $f \
                             --sample_ids $base \
                             -o filtered_q20/${base} -q 19 \
                             --barcode_type 'not-barcoded' --phred_offset=33
done
```

Chimera removal

```
wget https://drive5.com/uchime/gold.fa --no-check-certificate
mkdir chimeras

for f in $(find filtered_q20 -name "seqs.fna"); do
    base=${f#*/}
    base=${base%/*}
    identify_chimeric_seqs.py -m usearch61 -i $f -r gold.fa -o chimeras/$base
done

mkdir nonchim
for f in $(find filtered_q20 -name "seqs.fna"); do
    base=${f##*/}
    base=${base%/*}
    filter_fasta.py -f $f \
                    -o nonchim/${base}.fasta \
                    -s chimeras/${base}/chimeras.txt -n
done
```

Pick De Novo and Open Ref OTUs

```
mkdir otus_openref
mkdir otus_denovo

for f in $(find nonchim -name "*.fasta"); do
    base=${f##*/}
    base=${base%%.*}
    # Pick OTUs
    pick_open_reference_otus.py -i $f -o otus_openref/$base
    pick_de_novo_otus.py -i $f -o otus_denovo/$base
    # Export tables for R
```

```
    awk '{ print $1, NF-1 }' otus_openref/${base}/final_otu_map_mc2.txt > otus_openre
f/${base}_otus_openref_counts.txt
    awk '{ print $1, NF-1 }' otus_denovo/${base}/uclust_picked_otus/${base}_otus.txt
> otus_denovo/${base}_otus_denovo_counts.txt
done
```

##### Export tables for R

Run this from qiime1 folder to make a single OTU and single taxonomy table.

For De Novo OTU picking:

```
out="qiime1_diabimmune_otu_table.txt"
echo -e "sample_id\totu_id\tcount" > $out
for f in $(find otus_denovo -name "*_otus_denovo_counts.txt"); do
    sampleid=${f##*/}
    sampleid=${sampleid%%_*}
    awk -F" " -v sid=$sampleid '{ print sid"\t"sid"-"$1"\t"$2 }' $f >> $out
done

out="qiime1_diabimmune_tax_table.txt"
echo -e "sample_id\totu_id\ttaxonomy" > $out
for f in $(find otus_denovo -name "*_rep_set_tax_assignments.txt"); do
    sampleid=${f##*/}
    sampleid=${sampleid%%_*}
    awk -F "\t" -v sid=$sampleid '{ print sid"\t"sid"-"$1"\t"$2 }' $f >> $out
done
```

For Open Reference OTU picking:

```
out="qiime1_diabimmune_otu_table_openref.txt"
echo -e "sample_id\totu_id\tcount" > $out
for f in $(find otus_openref -name "*_otus_openref_counts.txt"); do
    sampleid=${f##*/}
    sampleid=${sampleid%%_*}
    awk -F" " -v sid=$sampleid '{
        # if (substr($1,1,3) == "New") { print sid"\t"sid"-"$1"\t"$2 }
        # else { print sid"\t"$1"\t"$2 }
        print sid"\t"sid"-"$1"\t"$2
    }' $f >> $out
done

out="qiime1_diabimmune_tax_table_openref.txt"
echo -e "sample_id\totu_id\ttaxonomy" > $out
for f in $(find otus_openref -name "rep_set_tax_assignments.txt"); do
    sampleid=${f#*/}
    sampleid=${sampleid%%/*};
        awk -F "\t" -v sid=$sampleid '{
        # if (substr($1,1,3) == "New") { print sid"\t"sid"-"$1"\t"$2 }
        # else { print sid"\t"$1"\t"$2 }
        print sid"\t"sid"-"$1"\t"$2
    }' $f >> $out
done
```
